# Thickness of deep layers in the fusiform face area predicts face recognition

**DOI:** 10.1101/788216

**Authors:** Rankin W. McGugin, Allen T. Newton, Benjamin Tamber-Rosenau, Andrew Tomarken, Isabel Gauthier

## Abstract

People with superior face recognition have relatively thin cortex in face-selective brain areas, while those with superior vehicle recognition have relatively thick cortex in the same areas. We suggest that these opposite correlations reflect distinct mechanisms influencing cortical thickness (CT) for abilities acquired at different points in development. We explore a new prediction regarding the specificity of these effects through the depth of the cortex: that face recognition selectively and negatively correlates with thickness of the deepest laminar subdivision in face-selective areas. With ultra-high resolution MRI at 7T, we estimated the thickness of three laminar subdivisions, which we term MR layers, in the right fusiform face area (rFFA) in 14 adult male humans. Face recognition was negatively associated with the thickness of deep MR layers, while vehicle recognition was positively related to the thickness of all layers. Regression model comparisons provided overwhelming support for a model specifying that the magnitude of the association between face recognition and CT differs across MR layers (deep vs. superficial/middle) while the magnitude of the association between vehicle recognition and CT is invariant across layers. The total CT of rFFA accounted for 69% of the variance in face recognition, and thickness of the deep layer alone accounted for 84% of this variance. Our findings demonstrate the functional validity of MR laminar estimates in FFA. Studying the structural basis of individual differences for multiple abilities in the same cortical area can reveal effects of distinct mechanisms that are not apparent when studying average variation or development.

**Significance Statement:** Face and object recognition vary in the normal population and are only modestly related to each other. The recognition of faces and vehicles are both positively related to neural responses in the fusiform face area (FFA), but show different relations to the cortical thickness of FFA. Here, we use very high-resolution MRI, and find that face recognition ability (a skill acquired early in life) is negatively correlated with thickness of FFA’s deepest MR-defined layers, whereas recognition of vehicles (a skill acquired later in life) is positively related to thickness at of all cortical layers. Our methods can be used in the future to characterize sources of variability in human abilities and relate them to distinct mechanisms of neural plasticity.

## Introduction

Visual abilities depend on genetic and environmental influences on mechanisms of neural plasticity. Brain structure is a task-independent biomarker that, in any one area, can relate to several skills. Given skills acquired at different times, variability in structure may reflect different plasticity mechanisms. We focus on a case in which the cortical thickness (CT) of *the same brain area in the same people* predicts different abilities in opposite directions: Vehicle recognition (VR) was associated with a relatively thicker FFA, whereas face recognition (FR) was associated with a relatively thinner FFA (McGugin, Van Gulick, & Gauthier, 2016). We used ultra-high resolution imaging to test hypotheses about the structural underpinnings of these behavioral abilities.

When first reporting this surprising pattern of results in FFA, we highlighted the earlier acquisition for FR compared to VR (McGugin et al., 2016). Three months-old babies see faces for 25% of their waking time, an amount reduced to 10% by 12 months (Fausey, Jayaraman, & Smith, 2016; Sugden & Moulson, 2018) and early face exposure has been found to be critical to the development of face-selectivity (Arcaro, Schade, Vincent, Ponce, & Livingstone, 2017). When infants learn to individuate objects such as vehicles is less clear, but in their first, infants’ visual diet includes a small set of repeating objects (e.g., cups and bowls) (Clerkin, Hart, Rehg, Yu, & Smith, 2016). Therefore, while face selectivity continues to develop during adolescence (Gomez et al., 2017; Scherf, Thomas, Doyle, & Behrmann, 2014) and FR continues to improve in adults (Germine, Duchaine, & Nakayama, 2011), it appears reasonable to assume FR starts earlier than when most people start to individuate vehicle models.

Learning to identify faces earlier than vehicles may influence their structural correlates in adult brains, since early development involves the pruning of connections. The amygdala is critical to the development of FR (Schultz, 2005) and is connected to the occipital lobe via the inferior longitudinal fasciculus (ILF). Structural properties of the ILF relate to face recognition (Gomez et al., 2015). We and others (Bi et al., 2014; McGugin et al., 2016) suggested that greater pruning of the infant ILF would result in more efficient connectivity and better FR in adults. Research on the connectivity between the inferior temporal (IT) areas TE and TEO (the monkey analog of the FFA) and the limbic system has revealed projections in the infant monkey that are pruned in the adult (Webster, Ungerleider, & Bachevalier, 1991). Such connections from IT to limbic areas are labeled predominantly in infragranual (deep) layers of IT (Stefanacci & Amaral, 2000). Thus, if this variability in the pruning of the ILF connections from IT to the amygdala is relevant to FR in humans, it should be most reflected in the thickness of deep layers in FFA.

A model instantiating differential early development would predict that the negative correlation between FR and CT will be stronger in deep layers than other layers. In contrast, we expect the positive correlation between VR and variability in FFA’s CT to be supported by mechanims that support learning-dependent structural plasticity in adults (Wenger et al., 2012). These mechanisms are not well understood but are proposed to include forces such as gliogenesis and angiogenesis that accompany synaptogenesis (Zatorre, Fields, & Johansen-berg, 2012). We have no reason to expect these mechanisms to be confined to deep layers (but acknowledge no evidence against laminar specificity).

We compare two sets of predictions. One specifies that the pattern of associations within each layer is the same as that observed for total CT (McGugin et al., 2016). Specifically, CT for each layer would be negatively associated with variability in FR but positively associated with variability in VR. We tested the most parsimonious version of this perspective (Model 1, Fig. 1) by specifying associations invariant across layers. In contrast, Model 2, the differential early development model, posits a selective linkage between the deep MR layer and variability in FR.

**Figure 1.**
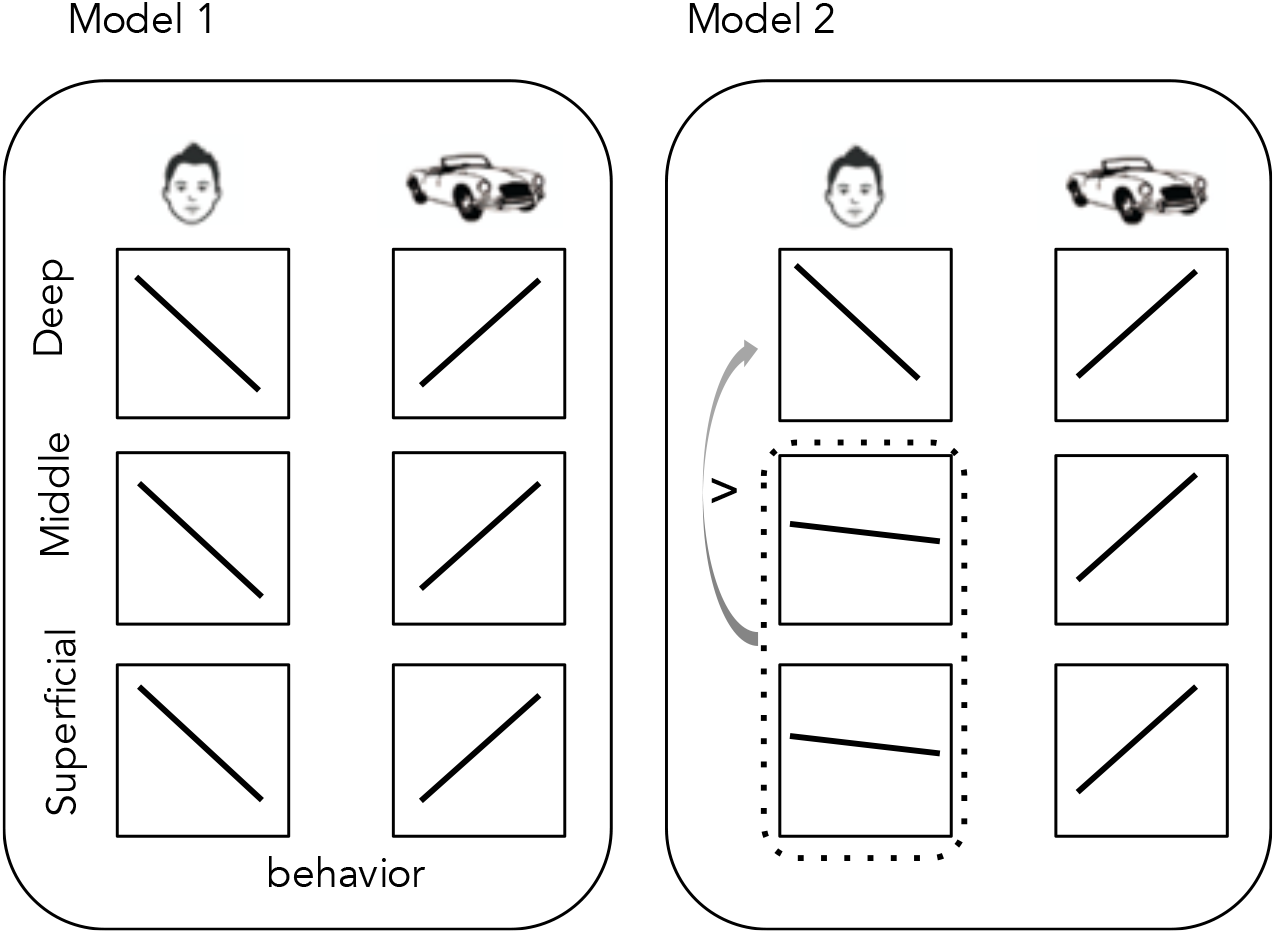
Composite Models tested. Model 1 specified that correlations and regression coefficients between face recognition and cortical thickness are negative in sign and equal across MR cortical layers and that correlations and regression coefficients between vehicle recognition and cortical thickness are positive in sign and equal across layers. Model 2 specified a layer-specific pattern for the associations between face recognition and cortical thickness across layers, with the deep MR layer having a stronger, more negative pattern of association than the superficial and middle MR layers. Like Model 1, Model 2 specified an invariant of association between vehicle recognition and CT across layers. As described in the text, we also tested three additional substantive models positing different patterns of association.

## Materials and Methods

### Subjects

14 right-handed men aged 19 to 29 (mean=22.46, s.d.=3.60) were recruited from the Vanderbilt University community based on previous participation. Only men were recruited to limit variability in brain anatomy. Our sample size was limited by the short period of time between when we developed scan protocols with sufficient resolution and before a major scanner upgrade. We recruited individuals with experience in the scanner, and who, as a group, offered a wide range of performance with faces. One subject was excluded due to excessive head motion. Written informed consent was obtained from all subjects in accordance with guidelines of the Vanderbilt University Institutional Review Board and Vanderbilt University Medical Center. All subjects reported normal or corrected-to-normal visual acuity, and received monetary compensation for participation.

### Behavioral Testing

Subjects performed a battery of matching and recognition memory tests outside of the scanner. First, they completed the extended Cambridge Face Memory Test (CFMT+; Russell, Duchaine, & Nakayama, 2009) in which they studied several views of unfamiliar male faces and were tested in forced-choice trials of three faces on which they selected the face that matched a studied face. Second, subjects completed various sections of the Vanderbilt Expertise Test (VET; McGugin, Richler, Herzmann, Speegle, & Gauthier, 2012). This included 2 face tests (VET-Males, VET-Females), 2 vehicle tests (VET-Car, VET-Plane), and 2 animal tests (VET-Bird, VET-Butterfly). In each case, subjects studied six identities, models or species for as long as needed, followed by 3-alternative forced choice test trials in which they indicated which item was studied. Third, subjects completed the Vanderbilt Face Matching Test (VFMT; Sunday, Lee, & Gauthier, 2018). On each trial, subjects studied 2 faces for 4000 ms, then were shown a 3-alternative forced choice and indicated which face was a new image of a face studied on that trial. Matching faces were different images from the original presentation. Fourth, subjects completed sequential matching tests using cars, planes, birds and butterflies (McGugin, Gatenby, Gore, & Gauthier, 2012). On each trial, an image appeared for 1000 ms, followed by a 500-ms mask, then a second image. Subjects judged if the two images showed cars/planes of the same make and model regardless of year, or birds/butterflies of the same species. See Table 1 for mean test performance, reliability of each test, and consistency across tests for each category. Aggregate Face, Vehicle and Animal indices were calculated based in each case on standardized performance for the four relevant tests.

**Table 1.** Behavioral results across twelve independent tests. For one subject, performance on VET-Male was near chance and an outlier relative to other subjects and his own other scores, so this data point was dropped. The ICC4 (with 95% HPD interval) for the aggregate of Face tests was .89 (.77,.96), for the aggregate Vehicle tests, .83 (.66,.95) and for the aggregate Animal tests, .85 (.70,.96).

|  |  | Mean (std dev) | Cronbach's Alpha |
| --- | --- | --- | --- |
| Faces | CFMT+ | 0.65 (0.13) | 0.88 |
|  | VET-Male (Acc) | 0.83 (0.17) | 0.78 |
|  | VET-Female (Acc) | 0.83 (0.12) | 0.86 |
|  | VFMT (Acc) | 0.6 (0.07) | 0.86 |
| Vehicles | VET-Car (Acc) | 0.73 (0.18) | 0.93 |
|  | VET-Plane (Acc) | 0.76 (0.12) | 0.93 |
|  | Matching-Car (d') | 1.79 (1.00) | 0.75 |
|  | Matching-Plane (d') | 1.57 (0.77) | 0.86 |
| Animals | VET-Bird (Acc) | 0.68 (0.17) | 0.80 |
|  | VET-Butterfly (Acc) | 0.56 (0.17) | 0.83 |
|  | Matching-Bird (d') | 1.46 (0.39) | 0.92 |
|  | Matching-Butterfly (d') | 1.65 (0.52) | 0.82 |

### MRI acquisition

All subjects were imaged using a Philips Achieva whole-body 7T MRI scanner with a quadrature, head only transmit coil and a 32 channel receive coil array. Imaging proceeded in three stages: whole brain structural imaging, functional localization, and ultra-high resolution susceptibility-weighted imaging (SWI).

### Whole brain structural imaging acquisition

Whole brain structural imaging consisted of a sagittal whole brain 3D T1 weighted MPRAGE image gathered at 1 mm isotropic resolution and resampled into axial and coronal orientations for accurate localization of SWI acquisitions. This was used to identify anatomic landmarks important to the planning of the subsequent ultra-high resolution SWI acquisitions.

### Functional MRI acquisition, stimuli, design and analyses

Functional localization of the fusiform face area (FFA) was accomplished via isotropic whole brain fMRI data. Images were acquired with minimal distortion using a multishot 3D gradient echo PRESTO EPI pulse sequence (Versluis et al., 2010). TR/TE=21.93/28.13ms, flip=12°, FOV=211.4mm, number of slices=30, voxel size=2.5mm isotropic, EPI factor=11, water/fat shift=6.705pix, number of dynamics=160, time/dynamic=2.02s).

### Experimental Design and Statistical Analysis

We presented all stimuli with an Apple Macintosh computer running Matlab 2014a (MathWorks, Natick, MA) using Psychophysics Toolbox (Brainard, 1997). Stimuli were displayed on a rear-projection screen using an Avotec projector. Immediately following the HR structural scan, each subject completed 1-2 runs (depending on time) of a functional localizer scan (160 dynamics/run). We used grayscale images (36 faces, 36 objects) in a 1-back detection task across 20 alternating blocks of face and object images. Each image was presented for 1 s, with 16 images per category block. The HR T1-weighted structural scans were transformed into the standard sagittal view with 1mm isometric voxels to facilitate coregistration of functional and anatomical datasets. Functional data were analyzed using Brain Voyager (www.brainvoyager.com) and in-house Matlab scripts (MathWorks, Natick, MA). Preprocessing in Brain Voyager included registration to the structural scan, slice scan time correction (cubic spline), 3D motion correction (sinc interpolation) and temporal filtering (high-pass criterion of 2 cycles per run) with linear trend removal. No spatial smoothing was applied. Regions of interest (ROIs) were defined using the Face>Object contrast from the face-localizer scan. We localized an ROI that responded more to faces than objects in the right middle fusiform gyrus (FFA2) of all subjects.

Right FFA2s were localized on the 3 mm^3^ (interpolated) statistical maps from the localizer scans as the peak face-selective voxel, then grown to incorporate contiguous 3-mm^3^ voxels passing statistical threshold within a 4-voxel or 108 mm^3^ fixed volume around the peak of face-selectivity. Solely for the purpose of comparing the location of our rFFA2s with those reported in the literature and demonstrating homology of the rFFA2 across subjects, the HR T1-weighted structural scans were normalized to Talairach space and functional data re-aligned to the Talairach-transformed images. Talairach coordinates for our ROIs (mm ± SD) are (X = 38.92 ± 3.86; Y = −47.54 ± 6.73; Z = −20.62 ± 3.01).

### SWI image acquisition, processing and averaging

A minimum of six ultra-high resolution T_2_* weighted images were acquired as separate and independent acquisitions in each subject using slice-selective Cartesian gradient echo acquisitions. Both real and imaginary images were recorded. Measurement of individual differences in CT presents additional challenges over characterizing average laminar profiles because of the various sources of measurement error that can confound inter-individual variability. Thus, care was taken to plan image acquisitions such that the frequency encoding direction was perpendicular to the ventral surface of the temporal lobe underneath the right FFA, to minimize differences in partial volume effects across layers from subject to subject. Prior to analysis, ultra-high resolution data were reconstructed at 0.1875 mm in-plane resolution.

SWI imaging parameters were as follows: FOV=240×180.194×21.9mm, yielding voxel dimensions of 0.194×0.194×1.00 mm, 20 slices, 0.1 mm gap, ‘shortest’ (878.8±8.29ms) repetition time, ‘shortest’ (27.5±0.31ms) echo time, 27.26 pix water/fat shift, 55 degree flip angle, flow compensation, 9 min 11 ± 5.2 sec total duration. Images were acquired with a 1D phase navigator to correct for phase errors arising from respiration during acquisition, as has been previously shown to be effective in very high resolution SWI imaging (Versluis et al., 2010).

Real and imaginary images were used to calculate magnitude and phase images, which were then processed to create susceptibility weighted scans (Haacke, Xu, Cheng, & Reichenbach, 2004). Within each subject, each of the six resulting SWI scans were co-registered using SPM12 (http://www.fil.ion.ucl.ac.uk/spm/) and averaged in MATLAB (https://www.mathworks.com). For those subjects that had more than six T_2_* weighted scans acquired, the optimal six were identified for analysis by calculating the normalized mutual information (NMI; Studholme, Hill, & Hawkes, 1999; Viola & Wells III, 1997; Wells, Viola, Atsumi, Nakajima, & Kikinis, 1996) between all pairs of acquired and co-registered SWI scans, and ranking each according to the sum of its pairwise NMI values. The six scans with the highest summed pairwise NMI were kept for further analysis. This ensured that the six scans most similar to each other, and likely containing the least substantial run-dependent artifacts or misalignments, were selected for the purpose of averaging in later stages of the analysis.

### Image registration

Standard-resolution functional localizer scans were registered to high-resolution (1mm isotropic) structural scans in BrainVoyager (Fig. 2A). Because we sought to avoid unnecessary interpolations of the ultra-high-resolution (UHR) images, all intensity analyses were performed directly on the UHR images. We registered the high-resolution structural image and the functionally-defined rFFA2 to the UHR slices. Specifically, high-resolution structural images and rFFA2 masks were converted from BrainVoyager’s proprietary image formats to NifTI-1 images using the NeuroElf toolbox for Matlab. Rigid-body transformation matrices of the high-resolution structural images to the UHR images were calculated via SPM’s *SPM_coreg()* coregistration function, and these matrices were used to reslice the rFFA2 masks into the UHR space. The 108 mm^3^ rFFA2 masks traversed 4 to 7 UHR slices depending on the subject (mean= 5.57, SD= .85). For each UHR slice, 16-bit TIFF-format two-dimensional slice images were saved using Matlab.

**Figure 2.**
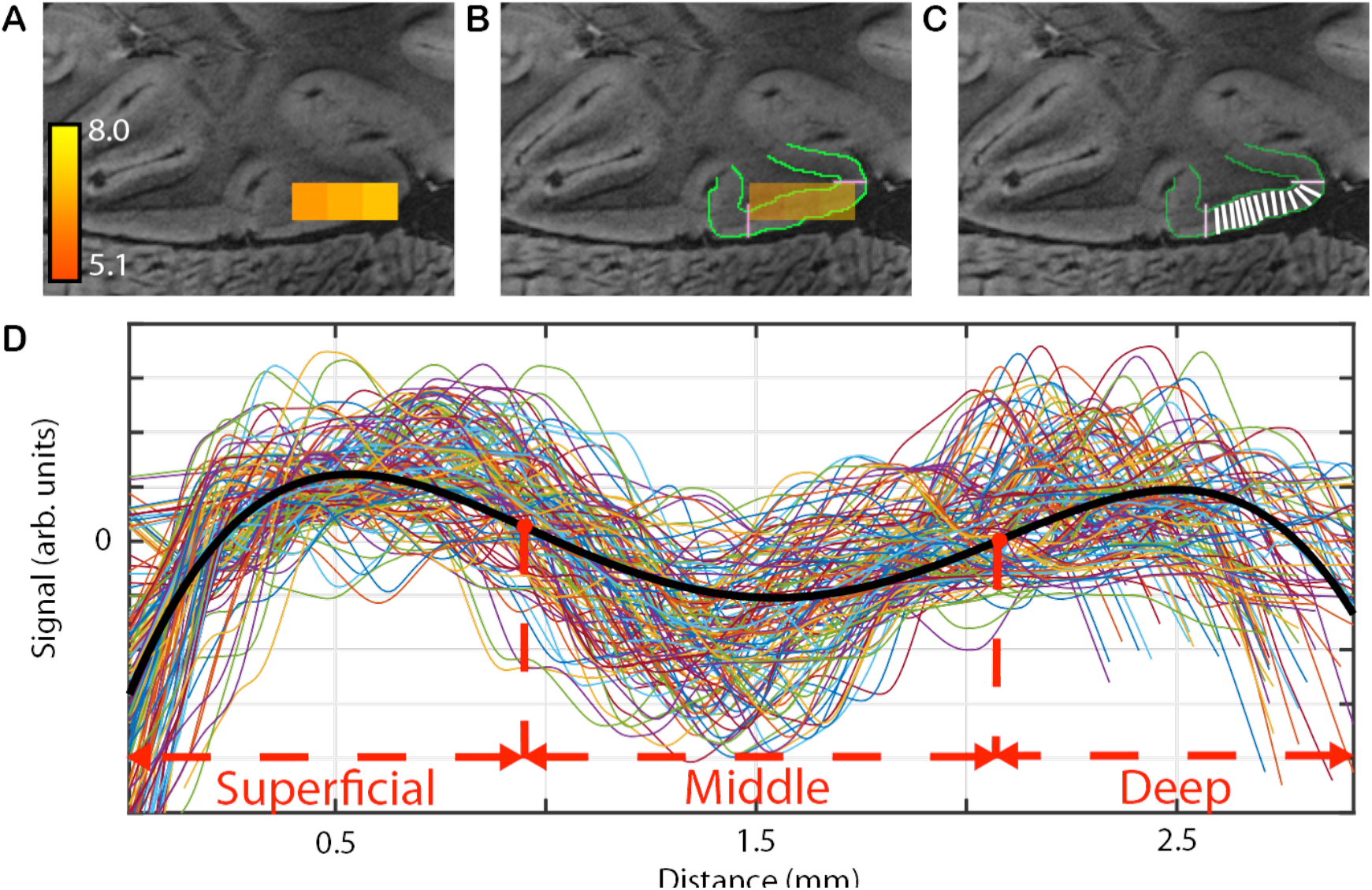
Trace acquisition, intensity sampling and layer identification. **(A)** rFFA2 was defined functionally with a Face > Object contrast and at a fixed size of 108 mm^3^ in each subject, then co-registered onto the ultra-high resolution slices. Color bar and values represent the face selectivity measured from the 3mm scan. **(B)** The edges of the rFFA2 were traced (magenta) over the underlying cortical mantle. Boundaries between grey matter (GM) and cerebrospinal fluid (CSF) (superficial layer boundary), and between GM and white matter (WM) (deep layer boundary), were manually drawn (green). **(C)** Traces were acquired as the series of lines originating at each superficial boundary voxel and terminating in each deep boundary voxel, and vice versa. Gray matter intensity was sampled across the length of the trace. **(D)** The line plot visualizes all traces shown in varying colors contributing to one slice of one subject’s rFFA2. A 4^th^-order polynomial (bold black curve) was fit to an average trace, with points of inflection defining the depths at which image intensity changed.

### Image Segmentation and Trace Identification

Using Adobe Photoshop, the gray matter/white matter (deep) and gray matter/CSF (superficial) cortical boundaries were manually drawn by two image analysts trained to recognize differences in brain structures but blind to the experimental goals (Fig. 2B). The inter-rater reliability of total rFFA2 thickness was very high (intraclass correlation, ICC=0.98). In addition to segmenting cortex from other tissue, blood vessels and image artifacts were manually marked as well. The resulting segmented data images were masked with the ROI images to create a series of ROI-specific cortical segmentation images. These segmented ROI images were next used to trace within-slice line segments through the thickness of cortex, along which we sampled the cortical signal intensity. Prior to tracing, each boundary was smoothed to avoid single-pixel concavities or convexities. Specifically, for each voxel that was part of the boundary, the three contiguous nearest neighbor boundary voxels in either direction (for a total of seven) were identified and a line segment fit to those voxels using linear regression. The set of such line segments was aggregated as the boundary was traversed, and skeletonized (to removed thickenings) and cleaned of spurs using the Matlab function *bwmorph()*.

Because of the convoluted shape of the cortical sheath, points on the deep and superficial boundaries of cortex do not have a one-to-one correspondence. Using custom Matlab code, we selected lines originating in each superficial boundary voxel and terminating in the nearest deep boundary voxel within that ROI in that slice (Fig. 2C). We conducted the same procedure in the reverse direction as well, originating in each deep boundary voxel and terminating in each superficial boundary voxel. The resulting traces were subject to five constraints: First, traces could not intersect cortical boundaries (aside from at trace origination or termination), to ensure that all samples were restricted to cortical tissue even in areas of high cortical convolution. Second, no trace could cross a previously-identified trace originating from the same boundary to avoid over-representing certain pieces of the cortical mantle. Third, the angle between the trace and boundary from which it originated could not be more than 20 degrees off perpendicular, to ensure that traces were approximately perpendicular to the cortical boundary. Fourth, any trace that was three or more standard deviations longer or shorter than the mean of all traces in a given slice was removed, to control for outlier paths. Fifth, no trace could cross an area marked for exclusion, such as a vein, since MRI intensity values are not valid indicators of gray matter signal in veins. For any trace that violated one of these constraints, a new trace was drawn from the originating voxel to the next-closest voxel in the other boundary.

### Intensity sampling and MR layer identification

First, average CT was defined in each ROI in each slice per subject as the average distance between the GM-CSF border and GM-WM border. Next, three cortical depths were identified by the points at which gray matter intensity trends changed (Fig. 2D). Specifically, to estimate these depths, we fit a 4th-order polynomial to the set of intensity samples from each ROI in each slice. Gray matter intensity was sampled at 500 equally-spaced points along the length of each trace. This ensured that voxel intensities were sampled even when a trace only crossed a corner of a voxel. The sampled intensities were linearly interpolated to 1/100th mm intervals. The resulting intensities were thresholded since pixels above or below a certain intensity are assumed to reflect partial-volume or whole-pixel capture of non-grey-matter. Specifically, any intensity values three or more standard deviations from the mean (over the entire set of trace intensities) were not included. Subsequently, intensity data were de-trended.

In all slices, inflection points were identified as the depths at which the second derivative of the polynomial was equal to zero. Inflection points outside the range of depths that fell within cortex were discarded and the remaining points were taken as estimates of the boundaries between cortical layers. Thus, we operationalized three MR cortical layers as the distances between (1) GM-CSF border and the first inflection point (superficial cortical depth), (2) first and second inflection points (middle cortical depth), and (3) second inflection point and the GM-WM border (deep cortical depth). We hypothesize that these MR layers are related to, although they are not a direct measure of, traditional cortical layers defined based on histological properties (Lifshits et al., 2018).

In 2 of 69 slices analyzed, the variation of intensity through the cortex was insufficient for accurately fitting a polynomial to the underlying data. These slices were omitted from further analyses.

### Transcortical microvasculature

Transcortical microvasculature was segmented and mapped using a contour based local minimization routine. Having manually identified the deep and superficial boundaries of the functionally defined rFFA2, intermediary contours parallel to the cortical surface were drawn throughout the cortical thickness. Transcortical vessels were segmented by mapping local minima along each contour, and the vascular area was quantified as the percent of voxels within the rFFA2 labeled as belonging to transcortical vessels. The measure for each subject was an average from at least 4 and up to 7 slices, with a weighted average ICC of .80.

## Results

### Reliability of CT measurements

#### Slices

We estimated the reliability of cortical measurements across slices (after centering them on the peak of face selectivity) using the correlation of thickness estimates for each slice relative to all the other slices in each subject. For total regional CT, which was based on manual tracings of high-contrast boundaries, correlations were higher than .8 for all slices. As a result, total CT measurements were averaged over all available slices. Inter-rater reliability of total CT was calculated using intraclass correlations for distance from GM-WM boundary to GM-CSF boundary, averaged across all slices for the two analysts.

Thickness measurements of the middle MR layer were based on relatively lower contrast boundaries identified using an automated algorithm (see below). Correlations reached .5 for anterior slices but slices that were posterior relative to the peak of face selectivity showed little to no evidence of reliability. Inspecting individual images, this proved to be because FFA2 ROIs were located closer to the posterior slices of the ultra-high resolution stack of 20 images for most subjects. Our measurements were made after co-registration of six ultra-high resolution scans to increase contrast, thus the final slices of the stack were more likely to show co-registration artifacts due to motion or respiration. We therefore used only the 2-4 reliable anterior slices for each subject and dropped the 2-3 most posterior and unreliable slices, for all measurements involving an internal laminar boundary (thickness estimates for deep, superficial and middle MR layers).

#### Average Thickness Measures

The white matter (WM)/grey matter (GM) boundary and the GM/cerebrospinal fluid (CSF) boundary were traced manually by two image analysts. Across subjects, the mean total CT of rFFA2 (distance from the WM/GM boundary to the GM/CSF boundary) was consistent with previous work. Between the two analysts blind to behavioral measurements, inter-rater reliability of total rFFA2 thickness was very high: intraclass correlation (ICC) = 0.98). MR-defined laminar subdivisions are defined by two internal boundaries identified using an automated procedure (see Methods). The representation of each MR laminar subdivision proportional to total CT was also consistent with previous work (Fig. 2D). Across subjects, the mean total CT of rFFA2 (distance from the WM/GM boundary to the GM/CSF boundary) was in the range reported in extrastriate cortex (Fischl & Dale, 2000): average rFFA2 GM thickness = 3.37 mm, with a range between 2.50 and 4.67 mm. Furthermore, the representation of each laminar subdivision proportional to total CT was consistent with MR microscopy in area V4 in nonhuman primates (Chen, Wang, Gore, & Roe, 2012): superficial MR layer (1.07 mm, range: 0.74 – 1.61 mm; 31%), middle MR layer (1.04 mm, range: 0.64 – 1.49 mm; 31%), deep MR layer (1.26 mm, range: 0.56 – 2.72 mm; 38%). As an index of consistency across subjects, ICCs for the three laminar subdivisions were: superficial, 0.77, middle, 0.73, deep, 0.80.

### Association between Behavioral Measures and Total rFFA2 CT

Each behavioral measure was computed as the average of the standardized values of four tests. ICCs were greater than .84 in all cases. We replicated previous results showing that total FFA CT is negatively correlated with face recognition, (r=−0.83 *p*<.001) and positively correlated with the recognition of vehicles (r=0.70, p<0.001, Table 2), with the two correlations significantly different (t=8.86, p<.001). Face and Vehicle performance were not significantly related (r=−.33, p=.27) and partial correlations with rFFA2 controlling for performance on the other categories were comparable in value and statistical significance to the zero-order *r*s (Table 2, Fig 3A). We replicated these correlational results using the univariate observed-imposed bootstrap (e.g., Beasley et al., 2007). Multiple regression indicated that both Face and Vehicle scores independently predicted rFFA2 thickness (both ps <= .001) and together accounted for 60% of the variance on adjusted R^2^.

**Table 2.** Pearson’s correlations between behavioral performance and cortical thickness and laminar depth measurements. Zero-order Pearson correlation coefficients are shown for Face and Vehicle categories correlated with total regional thickness and cortical depths operationalized by the 4th order polynomial and its 2nd order derivative. Partial-order Pearson correlation coefficients are given for each behavioral category regressing out the other two (including Animals), correlated with total regional and laminar cortical depths. Significant correlations bolded.

|  |  | Total Regional Thickness | Superficial Cortical Subdivision | Middle Cortical Subdivision | Deep Cortical Subdivision |
| --- | --- | --- | --- | --- | --- |
| Zero-Order Correlations | **Faces** | **-0.83** | -0.38 | -0.28 | **-0.76** |
|  | **Vehicles** | **0.70** | 0.44 | 0.48 | 0.45 |
| Partial Correlations | **Faces** | **-0.90** | -0.28 | -0.15 | **-0.73** |
|  | **Vehicles** | **0.83** | 0.37 | 0.43 | 0.33 |

**Figure 3.**
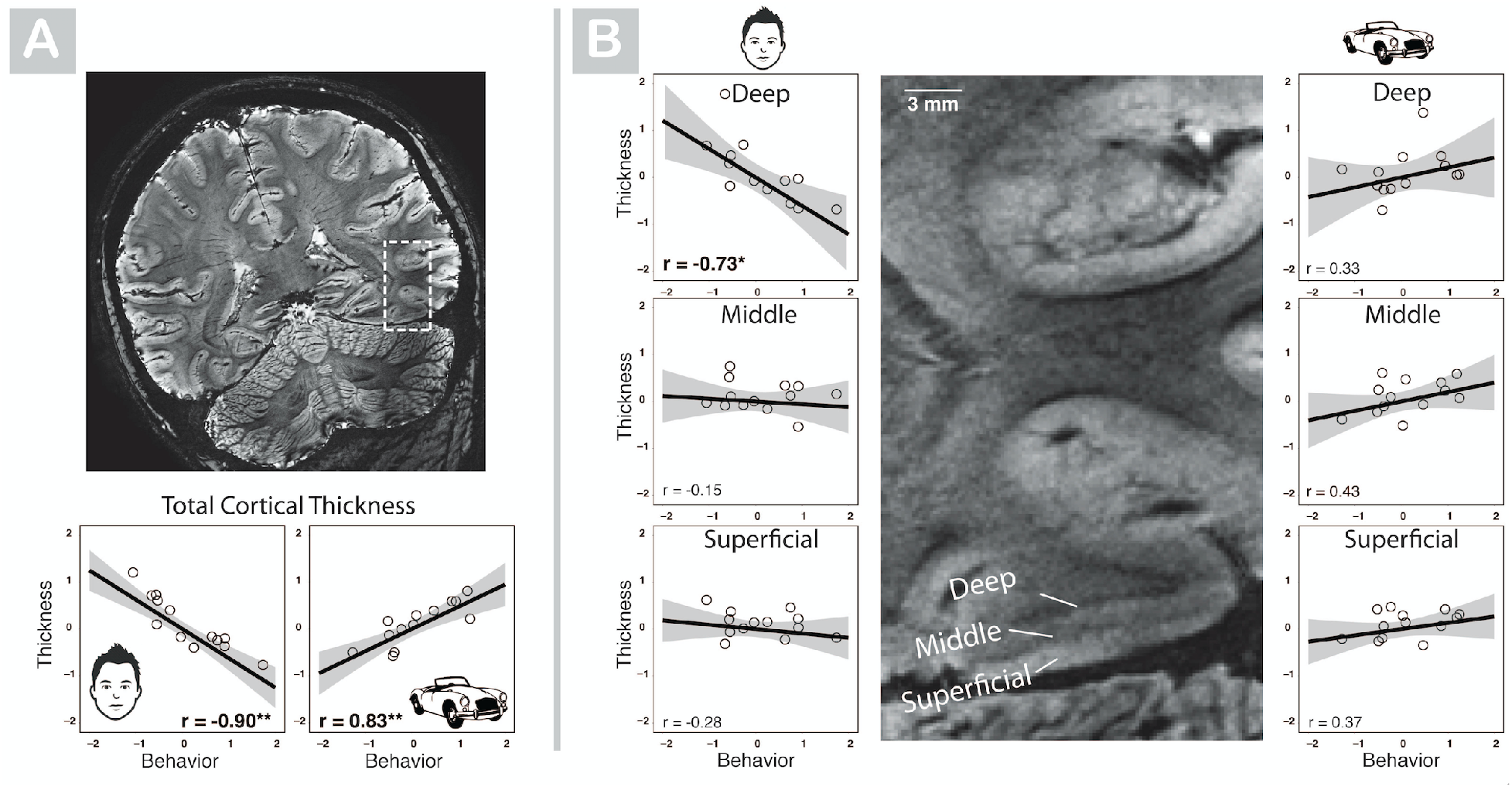
The relation between grey matter thickness of rFFA2, or its laminar subdivisions, and behavioral performance. Behavioral performance on the x-axes and cortical thickness and depth measurements on the y-axes show residualized values (see text for actual values in mm). The scatterplots on the left (with face icons) show partial correlations with Face recognition controlling for performance with Vehicles, while scatterplots on the right (with car icons) show partial correlations with Vehicles controlling for Face recognition. **(A)** The brain is tilted as a result of slices being aligned perpendicular to the rFFA2. Shown is one slice for one subject showing the average of six susceptibility-weighted scan acquisitions. The dashed box is centered on the functionally-defined rFFA2. Below the whole brain slice are scatterplots depicting the relation between total regional CT of rFFA2 and behavioral performance with Faces (left) and Vehicles (right). Error bars represent the 95% confidence intervals. **(B)** The central inset shows a magnified view from the box in (A), centered on rFFA2 for one slice in one subject, with the three laminar subdivisions perceivable to the naked eye. Scatterplots represent partial correlations between the thickness of Deep, Middle or Superficial MR layers as a function of face and vehicle recognition, controlling for performance with the other category. Error bars show the 95% confidence intervals.

As in prior work (McGugin et al., 2016), the correlation between overall CT and recognition of animals, which may not be acquired especially early or late, was in the same direction as face recognition although not significantly so (r=−0.12, p=0.34). Animal recognition also did not predict thickness of MR layers (superficial: r=0.15; middle: r=0.26; deep: r=−0.26). Our results cannot be explained by variability in vascular volume in FFA2 because such differences were not related to total CT (r=−.31), or behavioral performance (face, r=.11; vehicle, r=−.09, animal, r=.21), despite being reliable (ICC=.75).

### Association between Behavior Measures and Thickness of rFFA2 MR layers

#### Correlations

In accord with predictions of the differential early development model, face recognition was strongly negatively correlated with the thickness of deep MR layers of rFFA2 (r=−0.76, p=.002, Fig. 3B, Table 2). This relation is evident when examining subject-specific brain data listed as a function of face recognition performance (Fig. 4). There were no significant correlations between face recognition and thickness of superficial or middle MR layers (Fig. 3B, Table 2). In contrast, correlations between vehicle recognition and thickness of all three MR layers were consistently positive in sign (*r*s ranged from .44 to .48, two-tail *p*s ranging from .13 to .10), although not statistically significant. For the deep MR layer, the correlations of thickness with face recognition significantly differed from that with vehicle recognition (t=3.69, p<.005). These effects were replicated when bootstrapping was used.

**Figure 4.**
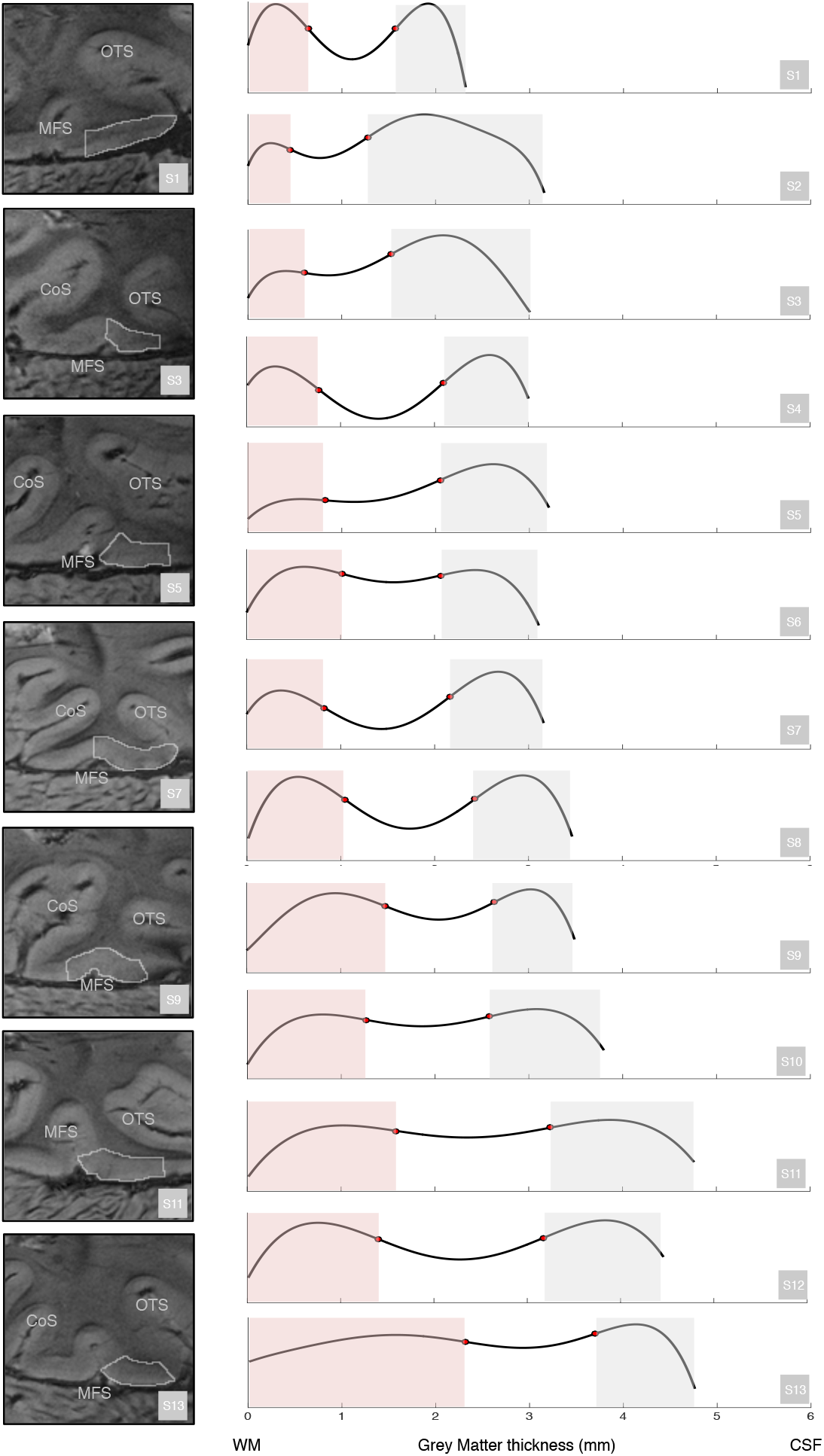
Plots of the 4^th^ order polynomial fit to the average rFFA2 intensity curve of the center-most slice for each subject. Subjects are rank-ordered from highest (S1) to lowest based on face recognition performance, regressing out performance with vehicles. Inflection points marking the transitions from deep MR layers (red shading) to middle MR layers (no shading) to superficial MR layers (grey shading) are marked with red dots. Axis labels: white matter (WM), cerebrospinal fluid (CSF). For every other subject, the centermost slice of rFFA2 is shown to the left. The spread of face-selective activation along the fusiform gyrus is outlined in grey. Labels: Occipital temporal sulcus (OTS), Collateral Sulcus (CoS), Middle Fusiform Sulcus (MFS). The rFFA2 most often fell on the lateral aspect of the fusiform gyrus, between MFS and OTS, corresponding to cytoarchitectonic FG4.

#### Regression Models Predicting Behavioral Measures From MR Layer-Specific CT

While correlations revealed bivariate relations among pairs of variables, they do not address the degree to which the combination of MR layer-specific CT measures predict individual differences in behavior. To address this question, we conducted multiple regression analyses in which CT in the superficial, middle, and deep MR layers predicted face recognition and vehicle recognition scores. According to adjusted R^2^, the combination of superficial, middle, and deep MR layer thickness accounted for 65% of the variance in face recognition (FR; overall regression F(3,9) =8.29, p =.006; unadjusted R^2^=.73). Both standard and bootstrapped confidence intervals indicated that the thickness of the deep layer was the only statistically significant predictor of face recognition. We also computed semi-partial correlations indicating the correlation between FR and each CT measure with the latter adjusted for the other CT measures. Only the semi-partial correlation for the deep MR layer (r = .72, p =.0125) was significantly greater than 0 (superficial r = −.29, p = .40; middle r = −.25, p = .45). Squared semi-partial correlations quantifying the proportion of variance uniquely contributed by each predictor indicated that the thickness of the deep MR layer accounted for slightly over half the total variance of FR (deep sr^2^ = .52, superficial and middle sr^2^ = .08 and .06 respectively).

The multiple regression analyses with vehicle recognition (VR) as the dependent variable had a different pattern of results. In this case, the overall regression model was significant (F(3,9) = 3.98, p < .05) although none of the regression coefficients for the three predictors were significant (ps ranged from .07 to .12). Although the latter result could be due to the relatively small sample size, this result also suggests that it was the shared variance among the three predictors rather than the unique variance of each predictor that accounted for effects on VR.

#### Regression Models Predicting CT of MR Layers from Behavior

Our most extensive analyses used a multiple regression framework to assess alternative models of the effects of face recognition and vehicle recognition on CT measures. We emphasized these analyses because they allowed us to test alternative hypotheses about the *complete pattern of associations* between the two behavioral measures and thickness of the three MR layers.

Our regression models used a repeated measures framework, with the 3 MR layer-specific CT measures per person serving as the repeated dependent measure. Predictors were layer, FR, and VR. We created two new variables that decomposed the overall effect of the layer factor into two orthogonal (i.e., uncorrelated) contrasts. As noted above, we hypothesized that the association between FR and thickness would be more strongly negative in the deep MR layer relative to the superficial and middle MR layers. To capture this difference, we gave one contrast variable (denoted L_DvSM_) a code of −1 for observations from the deep MR layer and codes of .5 for observations from the superficial and middle MR layers. The second contrast variable for layer (denoted L_SvM_) was orthogonal to the first and assigned a code of 0 to the deep layer, 1 to the superficial layer, and −1 to the middle layer. When both contrast variables are included in a model as a set, they estimate the overall (2 degree of freedom) main effect of MR layer.

We compared six different regression models that imposed different restrictions on the association between measures of the CT of MR layers (hereafter “layers” for short in the model descriptions) and FR and VR.

Model 0 (M_0_): Baseline Model.

In this model, the only predictor of CT levels was the overall main effect of layer:

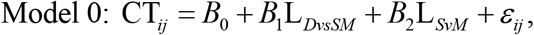

where i denotes subject (1 through 13), j denotes layer (1 through 3), and *ε* denotes the prediction error for the CT measure in the jth layer for the ith subject. We used M_0_ simply as a baseline comparison to verify that FR and VR measures, introduced in subsequent models, were significantly associated with CT.

Model 1(M_1_): Equal Coefficients Across Layers.

The next two models reflected our two primary alternative hypotheses. Model 1 predicted CT in a given layer from the two contrast variables for layer, FR, and VR:

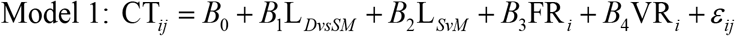

M_1_ is a “main effects only” model with no interaction terms. As a result it specifies that the regression coefficient denoting the effect of FR on CT (*B*_3_) is equal (i.e., invariant) across all three layers and the coefficient for the effect of VR on CT (*B*_4_) is also equal across layers (see Figure 1). This model is the most parsimonious extension of our previous finding using overall CT (McGugin et al., 2016).

Model 2 (M_2_): FR Differentially Predicts Deep vs Superficial/Middle MR Layer CT.

Model 2 tested the hypothesis that: (1) The association between FR and CT is more pronounced (i.e., more negative) in the deep layer than the middle and superficial layers; but (2) Layer does not moderate the relation between VR and CT measures. To embody this specification, an interaction (i.e., product) term between FR and the Deep vs. Superficial/Middle contrast variable (L_*DvsSM*_) was added to Model 1:

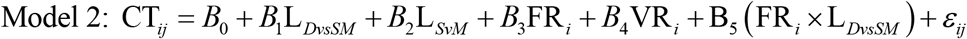

In model 2, three unique regression coefficients are sufficient to account for the linkages between FR and VR and CT across layers: the coefficient for the regression of CT on FR within the deep layer (equal to *B*_3_ − *B*_5_); the coefficient for the regression of CT on FR within the middle and superficial layers (equal to 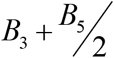); and, the single coefficient for the regression of CT on VR within each of the three layers (*B*_4_). Thus, this model specifies a constant effect of VR across all three layers and the hypothesized variable effect for FR.

The remaining models are designed to assess patterns of association other than those directly hypothesized in Models 1 and 2:

Model 3 (M_3_) extended Model 2 by adding an interaction term between the second Layer contrast variable (L_*SvM*_) and FR. This addition relaxes the assumption that the regression coefficients for FR in the middle and superficial layers were equal to one another. Thus, M_3_ goes beyond M_2_ by allowing all three regression coefficients for FR to differ from one another. Like M_2_, however it restricts all three regression coefficients for VR to be equal.

Model 4 (M_4_) added the full VR X Layer interaction to Model 1. This model specifies that the regression coefficients involving FR are all the same but allows for unique regression coefficients for VR in each layer.

Model 5 (M_5_) extended M_3_ and M_4_ by adding in both the overall FR X Layer interaction and the overall VR X Layer interaction. This model allows for 6 unique regression coefficients (FR/VR x Superficial/Middle/Deep) to capture the association between FR and VR and CT measures. Support for this model would imply that layer moderates the relation between *both* FR and VR and CT.

We conducted analyses using the gls function (Gałecki & Burzykowski, 2013) in the nlme package (Pinheiro et al., 2018) in R (R Core Team, 2018). For all models, we accounted for the dependence among the three observations for each subject by imposing an unstructured covariance matrix among the residuals. Maximum likelihood (ML) estimation was used (e.g., (Gałecki & Burzykowski, 2013). We evaluated the relative fit of the models using several indices. When models were nested, we used likelihood ratio tests to compare models. We also used three information indices to compare all models: The Akaike Information Criterion (AIC; Akaike, 1973), the small-sample corrected AIC (AICc; Sugiura, 1978), and the Bayesian Information Criterion (BIC; (Schwarz, 1978)). These indices reward model fit but penalize for model complexity, and thus are designed to prevent overfitting. Lower values indicate better fit. We also report −2 X the log likelihood (−2LL) value of each model computed at the maximum likelihood estimates. −2LL values are used both in the computation of likelihood ratio tests (for which the difference in the −2LL values of two models are distributed as chi-square variables with degrees of freedom equal to the difference in the number of parameters estimated by each model) and of information indices.

Table 3 shows the values of −2LL and the information indices for the six models. The values for M_0_ very clearly fit worse than all other models (e.g., all LR tests comparing other models to M_0_ ps < .005), thus indicating that FR and VR contributed significantly to the prediction of CT. Most importantly, Table 4 indicates that M_2_ (“FR Differentially Predicts Deep vs Superficial/Middle CT”) is clearly the best-fitting model. It has the lowest values for all three information indices. Likelihood ratio tests comparing M_2_ to other models yielded a pattern consistent with the information indices. M_2_ fit significantly better than M_1_ (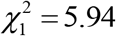, *p* =.02). Although M_2_ is a restricted version of M_3_ and M_5_, neither fit any better than M_2_ (M_2_ vs. M_3_ : 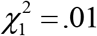, *p* >.90; M_2_ vs M_5_ : 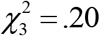, *p* >.97). These results indicate that M_2_ is the model that best combines fit to the data with parsimony. Table 5 shows the unstandardized and standardized regression coefficients for the regression of CT on FR and VR that are implied by M_2_. FR has a stronger (negative) effect on CT in the deep MR layer relative to the superficial and middle MR layers. This difference is statistically significant (t(33) = 2.53, p < .02).

**Table 3:** Fit Indices for Regression Models Predicting Cortical Thickness. -2LL =−2 * log likelihood of the model. This index is used in the computation of likelihood ratio tests comparing nested models and information indices. AIC = Akaike Information Criterion. AICc = AIC with small-sample correction. BIC = Bayesian Information Criterion. As lower values of information indices indicate better fit, all favor M2 (in bold). Bayes Factors compare M2 to each of the other models. All BFs indicate greater support for M2 than all other models.

| Model | Description | -2LL | AIC | AICc | BIC | M2/M#<br>Bayes Factor |
| --- | --- | --- | --- | --- | --- | --- |
| M<sub>0</sub> | Layer Only | 43.97 | 61.97 | 68.18 | 76.95 | 161.04 |
| M<sub>1</sub> | M<sub>0</sub> +<br>Main effects for FR and VR | 13.51 | 35.51 | 45.29 | 53.81 | 38.85 |
| M<sub>2</sub> | M<sub>1</sub> + L<sub>DvsSM</sub> | 7.56 | **31.56** | **43.56** | **51.53** | ----- |
| M<sub>3</sub> | M<sub>1</sub> + Layer x FR Interaction | 7.54 | 33.56 | 48.12 | 55.18 | 3.07 |
| M<sub>4</sub> | M<sub>1</sub> + Layer x VR Interaction | 12.52 | 38.52 | 53.09 | 60.15 | 139.52 |
| M<sub>5</sub> | M<sub>1</sub> +<br>Layer x FR + Layer x VR Interactions | 7.34 | 37.34 | 58.21 | 62.29 | 19.90 |

**Table 4:** Unstandardized and Standardized Regression Coefficients from Model 2. Model 2 is the best-fitting regression model that predicts that the Face Recognition is selectively associated with CT in the deep layer but that Vehicle Recognition has the same association with CT in each layer. Unstandardized regression coefficients are on top and standardized regression coefficients are below in parentheses. *= p < .001

| Layer | Face<br>Recognition | Vehicle<br>Recognition |
| --- | --- | --- |
| Superficial | -.07<br>(-.10) | .18*<br>(.27) |
| Middle | -.07<br>(-.10) | .18*<br>(.27) |
| Deep | -.62*<br>(-.57) | .18*<br>(.27) |

Although the fit indices discussed so far clearly indicate that M_2_ is the best-fitting model, they do not precisely quantify the strength of the evidence for M_2_ relative to the other hypotheses. For this reason, we additionally computed Bayes factors (BFs; (Jeffreys, 1961; Kass & Raftery, 1995)) using the BayesFactor package in R (Morey & Rouder, 2018). We specified the same six regression models. Instead of correlated residuals, we specified a random effect for subject to model the correlation among the CT values for each person. For all models, we used the Jeffreys-Zellner-Snow mixture of g-priors specification introduced by (Liang, Paulo, Molina, Clyde, & Berger, 2008) (see also Rouder & Morey, 2012). Because BFs can be sensitive to prior distributions, we used three alternative specifications of the scale parameter of the inverse gamma distribution that, in turn, is used to specify the variances of the prior distribution for the regression coefficients. All three specifications produced identical conclusions. Below, we report the results for the default “medium” values of the scaling parameters. For each pair of models, BFs were computed as the ratio of the marginal likelihood of the first model to the marginal likelihood of the second. Although we caution against rules of thumb, Jeffreys’ criteria specify that BFs in the range 1-3,3-10,10-30, 30 to 100, and > 100 indicate, respectively, weak, substantial, strong, very strong, and decisive support for the hypothesis linked to the numerator of the BF. Reciprocal values indicate degree of support for the model in the denominator.

An examination of the BFs yielded by the 15 pairings of models indicated that M_2_ was clearly the best-fitting model (i.e., for each of its 5 comparisons, M_2_ had the best fit). The final column in Table 3 displays the Bayes Factors comparing M_2_ to each of the other models. Perhaps the most important index here is the BF of 38.85 for the M_2_ /M_1_ comparison. This value indicates that the evidence in the data for M_2_ is over 30 times as strong as the evidence for M_1_ . All other comparisons yielded what Jeffrey’s termed “strong” support for M_2_ except that involving M_3_. This is not surprising given that both models are highly similar. Both allow for a difference in the association between FR and CT between the deep and the superficial layers, and both specify that the regression coefficients for VR are the same across layers. Unlike M_2_, however, M_3_ allows for differences between the FR coefficients in the superficial and middle layers. That the evidence favoring M_2_ is over three times stronger indicates that the superficial and middle layer coefficients can be constrained to be equal. Thus, the best-fitting model in both the regression and Bayesian analyses (M_2_) required only three unique regression coefficients to capture the relation between CT and FR and VR across the three MR layers: the coefficient denoting the effect of FR on CT in the deep MR layer, the coefficient denoting the effect of FR on CT in the superficial and middle MR layers, and the coefficient denoting the effect of VR on CT in all three MR layers.

## Discussion

We used susceptibility-weighted MRI in a 7T magnet to measure the thickness of deep, middle and superficial MR layers, distinctions that cannot be resolved with conventional MRI approaches at 3T or below. While new developments in MRI have allowed some visualization of intracortical layers, validation of their functional relevance has been reserved to primary visual cortex (Trampel, Ott, & Turner, 2011). We were constrained to a single region of interest due to a limited field of view and the fact that accurate measurement of laminar CT with non-isotropic voxels depends upon careful alignment of slices perpendicular to the cortex of a targeted brain area. We selected the rFFA2 based on earlier robust functional expertise effects for non-face objects (McGugin, Van Gulick, Tamber-Rosenau, Ross, & Gauthier, 2015).

We replicate and extend the finding that vehicle recognition is positively correlated with CT of FFA (McGugin et al., 2016), whereas face recognition is negatively correlated with FFA thickness (Bi et al., 2014; McGugin et al., 2016). This surprising pattern of correlations led us to postulate different mechanisms of plasticity behind these effects, with the mechanism relevant to face recognition operating earlier and selectively influencing the deep layers of rFFA2.

To measure the thickness of MR layers, we applied an automated algorithm to define borders between layers using signal intensity shifts along perpendicular traces through rFFA2 (Methods and Fig. 2). We postulate that the three most prominent intensity differences are related to known laminar divisions in the fusiform, corresponding to the densely-packed deep/infragranular layers (layers V-VI), middle/internal granular (layer IV), and superficial/supragranular (layers I-III). We successfully visualized laminar structure in rFFA2 and our measurements of the thickness of MR layers were reliable.

Although we compared several different models of the data, our primary focus was on two specific alternatives (see Fig. 1). The first (M_1_) specified that the pattern of correlations observed in total CT (negative for faces and positive for vehicles) is constant across all laminar subdivisions. In contrast, M_2_ specified that the association between individual differences in FR and the thickness of the deep MR layer was stronger (i.e., more negative) than the association between FR and thickness of the superficial and middle MR layers. Like M_1_, M_2_ specified that the association between VR and CT was invariant across layer. Regression analyses consistently favored M_2_ relative to M_1_ and to the other models that we specified, across multiple indices (likelihood ratio tests, three information indices, and Bayes Factors). Comparisons of regression coefficients yielded by M2 indicated that the association between FR and the deep MR layer was significantly stronger than the association between FR and the superficial and middle MR layers. The thickness of the deep MR layer accounted for a much higher proportion of variance in FR than the thickness of the other two MR layers. Indeed, when considered in isolation, CT of the deep MR layer accounted for 58% of the variance in FR, which is 84% of the variance accounted for by all the layers together (using total CT). Thus, the strong association between total CT and FR is largely, if not exclusively, due to the deep MR layer. In contrast, for VR there was no evidence that the relation was moderated by layer, and the clearly best-fitting regression model (M_2_) constrained the regression coefficients for VR to be equal across layers.

The sample size of the present study (N=13) was rather small. Accordingly, we focused on regression models that specified the overall pattern of association across layers and behavioral measures, to increase power and precision (e.g., see Hoijtink, 2011). In the present case, for example, while the individual correlations between VR and CT in the superficial, middle, and deep layers were not significantly greater than 0, our regression models yielded significant effects for VR on CT when these were constrained to be equal across layers. More generally, that our analyses indicated the clear superiority of M_2_ even given the relatively small sample size is an indication of its strength as a representation of the underlying pattern of associations.

These results are amongst the most structurally precise correlates of behavioral ability to date, and an important validation of the functional relevance of our in-vivo laminar measurements. Future development of these methods to a larger field of view should allow exploration of how these fine structural effects relate to correlations of face recognition ability on a much more distributed scale (Elbich & Scherf, 2017). Our results also represent perhaps the most striking difference to date between face and non-face recognition. Face recognition has often been deemed special relative to the recognition of other categories (Farah, Wilson, Drain, & Tanaka, 1998), but many behavioral hallmarks of face expertise, such as the inversion effect or holistic processing, are observed with non-face objects like cars in expert subjects (Bukach, Phillips, & Gauthier, 2010; Curby, Glazek, & Gauthier, 2009). Face and car recognition also show similar degrees of heritability (Shakeshaft & Plomin, 2015) and similar correlations with domain-general object recognition ability (Richler, Wilmer, & Gauthier, 2017). In fMRI studies, car recognition ability correlates positively with selective neural responses to images of cars in FFA (e.g., McGugin, Gatenby, et al., 2012), while face recognition ability also correlates positively with selective responses to faces in the same area (McGugin, Ryan, Tamber-Rosenau, & Gauthier, 2018). Despite these similarities, our results suggest that behaviors supported by common circuitry in a brain region may have developed at different times, with variability in structure as a footprint of the different acquisition history.

Our predictions were inspired by the idea that face recognition ability begins earlier than vehicle recognition and that a skill acquired during this period would be influenced by the pruning of connections from IT to limbic areas, though this is not the only possibility. We do not know whether the relevant period of influence on FFA’s structure occurs before about 5 years of age (by that age, children show many signs of the expertise adults display with faces (Jeffery et al., 2011)), when pruning may be more important, or in later childhood and adolescence, when myelination may dominate developmental changes (Natu et al., 2018). An alternative explanation is that greater myelination near FFA during childhood and adolescence is associated with better face recognition. The effects of pruning and myelination are difficult to distinguish, as myelination shifts the apparent gray-white matter boundary and results in thinner measurements of deep cortical layers (Sowell et al., 2004). Recent work has questioned the prevalent theory that pruning is the main force behind cortical thinning during development, with more evidence for myelination than for pruning after 5 years of age in the fusiform gyrus (Natu et al., 2018). Also unknown are the specific mechanisms of plasticity that would support structural effects of visual learning in adults, leading us to assume the null hypothesis of no laminar specificity of the positive correlation with CT observed for vehicles. Importantly, the mechanism driving the largest amount of average change in CT in an area may not be the same mechanism driving individual differences in CT.

It is unknown whether negative correlations between CT and performance will prove to be specific to deep MR layers for other kinds of abilities and other brain regions. It is also unknown which of the two patterns (positive correlation with total regional CT or negative correlation with thickness of deep MR layers) is more representative of the majority of object categories. Here, as in prior work at 3T (McGugin et al., 2016), there was no significant correlation with animal recognition but the trend was more similar to effects found with faces than vehicles (a negative correlation for thickness of deep MR layers). Future studies should explore how laminar CT in FFA relates to a more extensive set of visual abilities, and how this relation is affected by sex. Indeed, as of now, the cross-over interaction between CT and visual recognition abilities has only been described in men, yet there are sex differences in the recognition of living and non-living objects (McGugin, Richler, et al., 2012), and sex also moderates CT effects (Plessen, Hugdahl, Bansal, Hao, & Peterson, 2014).

By the time CT is measured in adults, effects in any given brain area will reflect a complex history of changes due to both experience- and non-experience-dependent developmental factors. This complicates interpreting relationships between behavior and local brain structure, including how structure underlies various neurological disorders. For example, both abnormal thinning of fusiform cortex over development (Raznahan et al., 2017) and lack of developmental improvements in face recognition (O’hearn, Schroer, Minshew, & Luna, 2010) have been reported in Autism Spectrum Disorders. The study of the laminar specificity of structural correlates of behavioral abilities may provide a useful window into these complex dynamics.

## Acknowledgments

We thank Carla Shatz for suggesting that different ages of acquisition should translate into differential effects on cortical layers. We thank Magen Speegle and Susan Benear for manual tracing on the grey-matter/white-matter boundaries, Anita Disney, Jesse Gomez, Thilo Womelsdorf and Wolfe Zinke for comments, Maarten Versluis and Ed Mojahed for sharing pulse sequence designs which were used as a basis for those developed here. R.W.M. and I.G. conceived the study. R.W.M and A.T.N. collected the primary data. A.T.N. developed the image acquisition techniques and average susceptibility-weighted image reconstruction techniques. R.W.M., A.T.N. and B.T.R. developed the methods for laminar analysis of susceptibility-weighted images. A.T. conducted reliability, bootstrapping, and Bayesian analyses. All authors coordinated second level analyses. R.W.M., A.T.N. and I.G. wrote the paper with input from the other authors. This work was supported by the NSF (SBE-0542013 and SMA-1640681) and the Vanderbilt Vision Research Center (P30-EY008126) and the Vanderbilt University Medical Center (1 S10RR023047 01).

